# Bursts of amino acid replacements in protein evolution

**DOI:** 10.1101/210211

**Authors:** Anastasia V. Stolyarova, Georgii A. Bazykin, Tatyana V. Neretina, Alexey S. Kondrashov

## Abstract

Evolution can occur both gradually and through alternating episodes of stasis and rapid changes. However, the prevalence and magnitude of fluctuations of the rate of evolution remains obscure. Detecting a rapid burst of changes requires a detailed record of past evolution, so that events that occurred within a short time interval can be identified. Here, we use the phylogenies of the Baikal Lake amphipods and of Catarrhini, which contain very short internal edges facilitating this task. We detect 6 radical bursts of evolution of individual proteins during such short time periods, each involving between 6 and 38 amino acid substitutions. These bursts were extremely unlikely to have occurred neutrally, and were apparently caused by positive selection. On average, in the course of a time interval required for one synonymous substitution per site, a protein undergoes a strong burst of rapid evolution with probability at least ~0.01.

## Introduction

(Non)uniformity of the rate of evolution is one of the oldest and most contentious issues in evolutionary biology. On the one hand, evolution often occurs gradually, so that its rate is approximately constant (Martin and Palumbi, 1993; Kumar and Hedges, 1998; dos Reis et al., 2012; Tamura et al., 2012). In particular, this is the case for selectively neutral segments of genomes, which evolve at rates equal to the corresponding mutation rates (“molecular clock”, Zuckerkandl and Pauling, 1965; Kimura and Ohta, 1974). Instances of gradual adaptive evolution are also known (e. g., Cooney et al., 2017). On the other hand, evolution may also occur mostly through short bursts of changes alternating with long periods of stasis (“punctuated gradualism” or “punctuated equilibrium”, Gould and Eldredge, 1993; Stanley, 1998). Examples of punctuated equilibrium are provided by the evolution of mammalian body weight (Mattila and Bokma, 2008), hominoid body size (Bokma, 2002), several morphological traits of rockfish (Ingram, 2011), intersexual signalling of cranes (Ø Mooers, Vamosi and Schluter, 1999), and many other data (Wolf et al., 2006; Hunt, 2007, 2008; Strotz and Allen, 2013; Bedford et al., 2014; Hunt, Hopkins and Lidgard, 2015; Voje, 2016). Clearly, both gradual and burst-like evolution does happen, but their relative importance remains controversial (Pagel, Venditti and Meade, 2006; Venditti and Pagel, 2008; Pennell, Harmon and Uyeda, 2014).

In order to detect short bursts of changes, one needs to be able to identify evolutionary events that occurred during a short interval of time. This is easy to accomplish if a very detailed paleontological record is available, such as those that exist for some marine invertebrates, e. g., Foraminifera (Malmgren, Berggren and Lohmann, 1983). However, such records are exceptions rather than the rule. Furthermore, paleontological data usually shed light only on the morphology of organisms and, thus, cannot reveal bursts of changes at the level of genomes. Fortunately, it may be possible to identify such bursts indirectly, through comparison of genomes of extant species, as long as their phylogenetic tree contains very short internal edges. We took advantage of two phylogenetic trees that meet this requirement, those of the Lake Baikal amphipods (Naumenko et al., 2017) and of Catarrhini (Rosenbloom et al., 2015, UCSC 100 vertebrates multiple alignment), and investigated short bursts in the evolution of their proteins.

## Materials and methods

### Dataset

We used two datasets. First, we considered transcriptomes-based clusters of orthologous genes (COGs) with exactly one ortholog represented in each species for five clades of the Lake Baikal amphipods (gammarids) (64 species and 3399 COGs in total), and the phylogeny based on them. The size of a clade varies from 6 to 24 species (Supplementary Fig. 1A). Each clade consists of closely related species, which makes it possible to reconstruct ancestral states with high confidence. Thus, all amino acid substitutions that constitute a burst can be reliably identified. The search for bursts was performed for each clade separately.

Second, we considered a multiple alignment of protein-coding genes from 11 primates species obtained from the 100 vertebrates genomes alignment of the UCSC Genome Browser together with the corresponding reconstructed phylogenetic tree (Supplementary Fig. 1B). In total, there were 17755 alignments of protein-coding genes of primates containing columns without gaps.

Only internal edges, i.e. segments of the phylogenetic tree ancestral to more than one species, were used in our analysis. For both datasets, we only considered internal edges of length < 0.005 dS units. We used codeml program of the PAML package (Yang, 2007) to reconstruct substitution histories of sequences and to estimate dN/dS values. Only gapless alignment columns were considered.

Presumptive functions of amphipod genes were inferred from blast2GO predictions (Naumenko et al., 2017) and from the genome annotation of a related species *Hyalella azteca* (BCM-HGSC); functions of primate genes were inferred the human genome annotation (hg38).

### Analysis

We assumed the neutral null model (dN=dS), which makes it possible to detect only the longest and the fastests bursts (see Discussion). We related dN of the gene on a particular edge of the phylogenetic tree to the length of this edge. Length edge was measured in the units of dS on the basis of all the available genes. We used this approach, instead of considering the dS value of only the gene that underwent a burst, because it is impossible to estimate the dS for an individual gene on a short edge with precision. For example, the expected number of synonymous substitutions in a gene encoding a 200 amino acid long protein on the edge of length 0.005 dS is only ~1. For a particular edge, the p-value associated with a nonsynonymous burst was calculated for each gene as the probability of observing this many or more nonsynonymous substitutions in a Poisson distribution with the parameter equal to the edge length (dS) (or, for bursts spanning multiple edges in a row, the sum of their lengths) multiplied by the number of nonsynonymous sites in the gene (estimated according to Nei and Gojobori, 1986). The obtained p-values were adjusted for multiple comparisons using the Benjamini-Hochberg method.

As a control, we also searched for statistically significant bursts of synonymous substitutions on the same set of short edges using an analogous approach.

### Filtering

The primary set of putative bursts with Benjamini-Hochberg adjusted p-values <0.05 was filtered as follows.

First, alignments containing >50% of columns with gaps were excluded. Second, to ensure the precise phylogenetic positioning of all substitutions that constituted a burst, we excluded sites with codeml posterior probabilities for the reconstructed ancestral variant < 0.8 and recalculated the statistics for this gene.

Third, to safeguard against contribution of anciently divergent paralogs or pseudogenes rather than orthologs to our findings, we applied additional filtering. We required that the dS value that characterized the edge of the putative burst obtained using the considered gene was lower than the dS value for this edge obtained using all genes (adjusted p-value > 0.001). Genes with substitutions in multiple repeated domains were excluded. Next, we required the absence of evidence for paralogs or duplications of the considered gene as follows. For each gene, we determined the pre-burst sequence as the reconstructed sequence of the phylogenetic node immediately ancestral to the burst-carrying edge(s), and the post-burst sequence as the reconstructed sequence of the phylogenetic node immediately descendant to it. For gammarids, we mapped raw transcriptomic reads of the considered gene from all species onto the pre-burst and post-burst sequences. If any reads from any of the species descendant to the edge of the provisional burst supported the pre-burst variant, or if any reads from any of the species not descendant to the edge of the provisional burst supported the post-burst variant, this gene was discarded. For primates, we aligned the pre-burst and post-burst sequences onto the assembled genomes of this gene from all species, and proceeded analogously. For primates, we also required that the genomic position of each burst-carrying gene was conserved in all primates genomes.

Finally, the burst-containing alignments that survived these filters were curated manually. If most substitutions constituting a burst fell into regions of poor alignment or were located in the very beginning or the very end of the gene, the corresponding putative burst was discarded.

## Results

We searched for bursts of amino acid substitutions (“bursts”) within internal edges of phylogenetic trees that are shorter than 0.005 dS. Suitable edges are present in 5 clades of the phylogenetic tree of gammarids from the Lake Baikal (Naumenko et al., 2017): *Eulimnogammarus* and related genera (18 edges), *Pallasea* and related genera (10), Hyallelopsis (3), Acanthogammaridae s. str. (7), and Micruropidae (4); as well as within the Catarrini clade (3 edges) of the tree of vertebrates (Supplementary Fig. 1) (Rosenbloom *et al.*, 2015). A burst consists of several amino acid substitutions which occurred in a protein within such an edge or, perhaps, within several successive edges of combined length below 0.005 dS.

In gammarids, 5 statistically significant bursts occurred in 5 proteins within 2 clades. 3 of them occurred over the time period corresponding to a short individual edge of the phylogeny, while the remaining 2 spanned two very short adjacent edges. In Catarrhini, there was 1 significant burst (Table 1). Each burst consisted of between 6 and 38 amino acid substitutions, or between 6 and 39 nonsynonymous substitutions (as some amino acid sites underwent multiple nonsynonymous substitutions), scattered throughout the protein (Supplementary Fig. 2). All edges that harbor bursts have 100% bootstrap support. Genes that harbored bursts are enriched in mitochondrial proteins: they constitute 3 of the 5 such genes, although only 14% of the initial set of COGs are annotated as components of mitochondria (binomial test, p-value = 0.02); Supplementary Fig. 3) (Naumenko *et al.*, 2017). No significant synonymous bursts were observed.

**Table 1.** Bursts of evolution in proteins of the Baikal gammarids and Catarrhini.

| clade | edge number | edge length (dS) | gene name (predicted) | description of the protein | overall dN/dS for the gene (excluding the edge with the burst) | substitutions during burst |  | adjusted p-value |
| --- | --- | --- | --- | --- | --- | --- | --- | --- |
|  |  |  |  |  |  | nonsyn. | syn. |  |
| *Pallasea* and related genera | 1 | 0.0040 | *DNAJC11* | DnaJ-like protein subfamily c member 11 | 0.41 | 39 | 1 | 1.01e-25 |
|  | 1 | 0.0040 | *MRPL22* | mitochondrial ribosomal protein L22 | 0.55 | 10 | 1 | 0.012 |
|  | 2+3 | 0.0007 + 0.0008 | *NOP16* | nucleolar protein 16-like | 0.35 | 6 | 2 | 0.046 |
| *Eulimno-gammarus* and related genera | 4 | 0.0011 | *MRPS25* | mitochondrial ribosomal protein S25 | 0.31 | 6.5 | 0.5 | 0.0011 |
|  | 5+6 | 0.0024 + 0.0010 | *AKR1* | aldo-keto reductase | 0.57 | 10 | 1 | 0.032 |
| primates | 7 | 0.0043 | *PKR* | interferon-induced protein kinase R | 1.27 | 18 | 2 | 0.0010 |

The most remarkable burst involving 39 nonsynonymous substitutions occurred in the mitochondrial chaperone gene (DNAJC11) on an edge of length 0.004 dS in the *Pallasea* clade (Fig. 1). This edge also harbored another burst in a mitochondrial protein (L22 ribosome protein) (Table 1). Bursts that are confined to one edge of the phylogenetic tree do not extend to preceding and/or successive edges (p-values for such edges > 0.27) (Supplementary Fig. 4). Hence, the characteristic duration of a burst is short, ~10^−3^ dS. The overall rate of evolution of some burst-carrying genes was somewhat higher than the average (Table 1, Fig. 2). Still, after multiple testing correction, there remains no genes with more than one statistically significant burst (p-values > 0.018, adjusted p-values = 1).

**Fig. 1.**
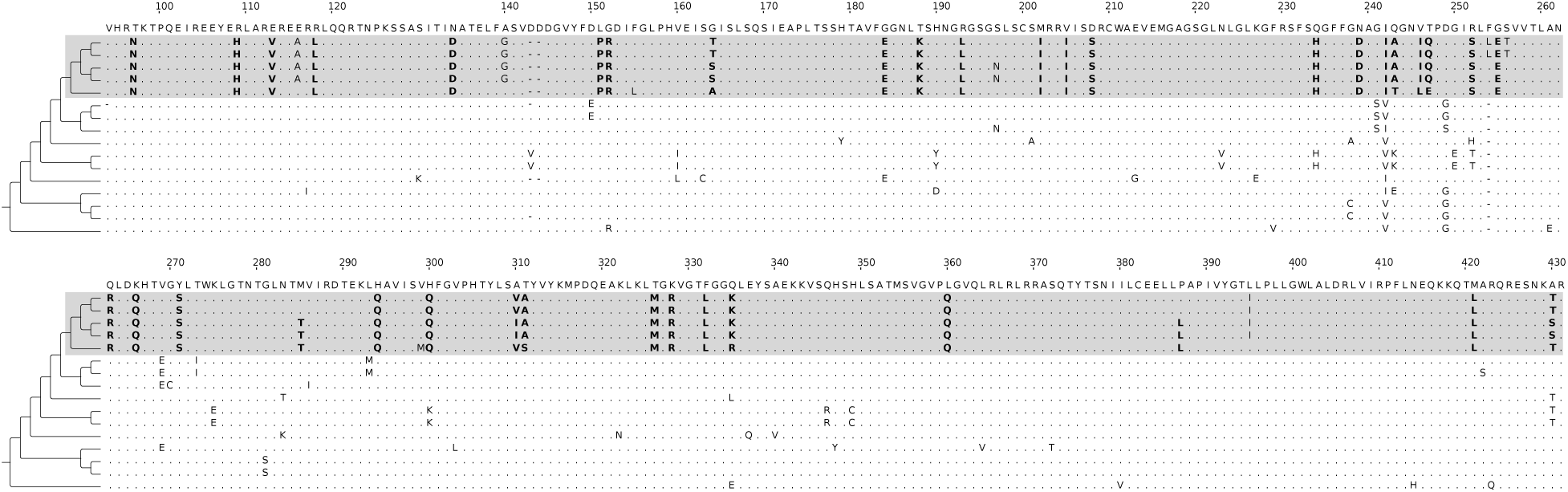
A fragment of the alignment of orthologous DNAJC11 genes of Pallasea gammarids, which contains 39 nonsynonymous substitutions on the internal edge 0.004 dS in length. Alleles derived in the adaptive burst are shown in bold. Total length of the alignment is 1677 nucleotides, or 1515 nucleotides without gaps.

**Fig. 2.**
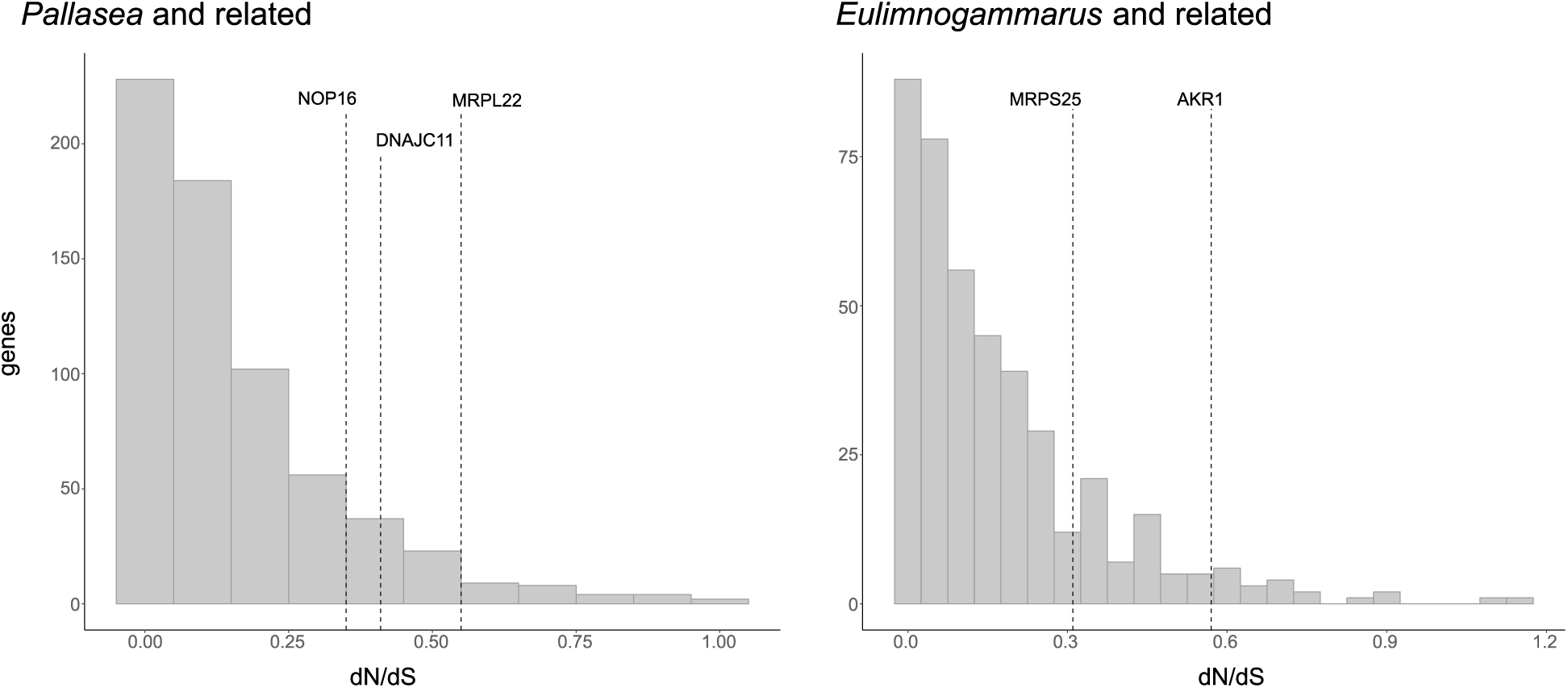
Distribution of dN/dS in genes of Baikal Amphipods in the clades carrying bursts. Genes with confirmed bursts are shown with dashed lines.

Phylogenetic tree of 11 species of Catarrhini has only three internal edges shorter than 0.005 dS (Supplementary Fig.1B), and only one statistical significant burst has been detected. This small number may be due to several reasons: Catarrhini species are more distant from each other, which results in longer phylogenetic edges and less confident ancestral state reconstruction; moreover, a larger initial dataset leads to a more substantial multiple testing correction.

The detected burst occurred on the internal edge ancestral to two macaques species. The gene with the burst encodes the PKR protein (also known as EIF2AK), which is the eukaryotic translation initiation factor 2 kinase activated during viral infection. As in gammarids, the substitutions are scattered along the sequence (Fig. 3). Primate PKR contains two dsRNA binding motifs (DRBMs 1 and 2) and C-terminal catalytic kinase domain. The kinase domain carries 14 amino acid substitutions on the selected edge, and DRBM1 the remaining 4. Most substitutions lie in αD, αG and αH helices or nearby, which have been shown to be enriched in positively selected sites (Rothenburg et al., 2009). αG helix and specifically positions with amino acid substitutions on the selected edge are involved in PKR interaction with eIF2α (Krishna et al., 2016); however, there is no evidence of bursts of evolution in eIF2α gene on the same edge.

**Fig. 3.**
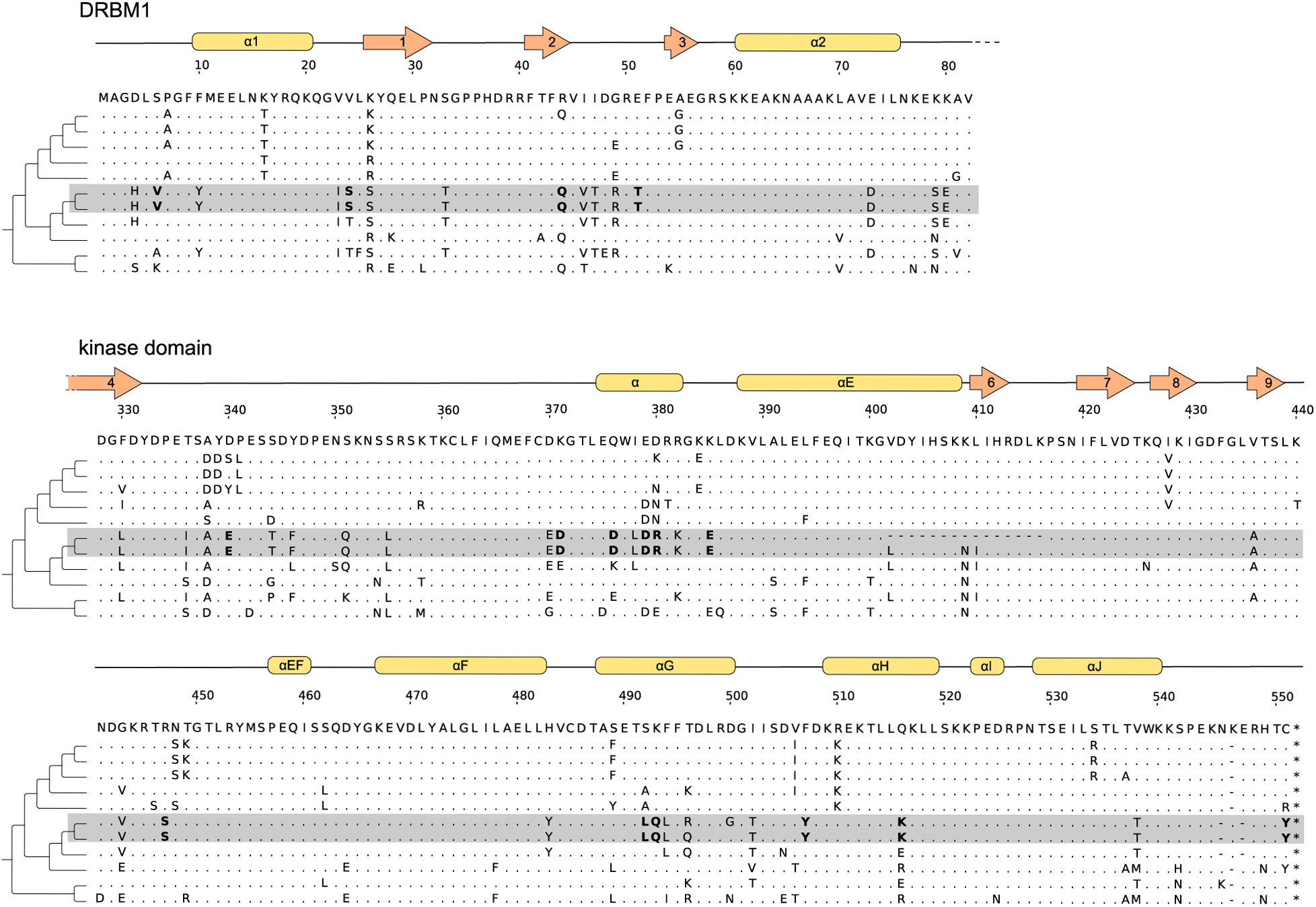
Alignment of PKR genes of Catarrhini (fragments) containing 18 nonsynonymous substitutions on the internal edge ancestral to Macaca mulatta and Macaca fascicularis. The majority of substitutions (14) occurred in the kinase domain of the protein, the other fall at the dsRNA binding motif (DRBM1). Alleles derived in the adaptive burst are shown in bold.

## Discussion

Allele replacements driven by positive selection are the fundamental genetic mechanism of adaptive evolution. These replacements can occur independently of each other or be correlated (Bazykin et al., 2004). *A priori*, there is a continuum of possibilities, from fully independent individual substitutions to bursts of adaptive evolution, each consisting of multiple substitutions that occurred over a short period of time. We searched for such bursts within individual proteins, taking advantage of two phylogenetic trees, of the lake Baikal gammarids (Naumenko *et al.*, 2017) and of Catarrhini (Rosenbloom et al., 2015).

Using only internal edges is essential because in this case the derived sequence is observed in more than one species, so that rare sequencing and alignment errors would not lead to false discovery of bursts. Our criteria for detection of bursts were rather stringent: conservative filtering of alignments, a neutral null model (dN = dS), and multiple testing correction. Unfortunately, it is hard to define the correct null model which takes into account all possible features of protein evolution, for example, those resulting from non-uniformity of the mutation rate along the genomes. Thus, our p-values should be viewed with caution. Still, the 6 bursts that we have found are likely to be “real”, in the sense of being caused by simultaneous or near-simultaneous action of positive selection at multiple sites within a protein.

Mutations that initiated amino acid substitutions that together constitute a burst are extremely unlikely to appear simultaneously as parts of one complex mutational event. Thus, a burst is likely to involve substitutions that were not precisely synchronous. In other words, a burst lasts longer than a substitution. Still, the bursts that we detected are quite short, at the evolutionary time scale. Indeed, four bursts were confined to just one internal edge shorter than 0.005 dS. The remaining two bursts, involving 6 and 9 amino acid substitutions, each occurred on two successive internal edges, of lengths 0.0007 and 0.0008, and 0.0024 and 0.0010 dS, suggesting that ~3-4 nonsynonymous substitutions occurred per 0.001 dS of evolutionary time. Among the substitutions involved in such composite bursts, neither occured on both edges after a cladogenesis (multiple non-synonymous substitutions did not occur on edges leading to *Eulimnogammarus testaceus* or *Pallasea cancellus*).

Of course, the two phylogenetic trees which we studied almost certainly contained other bursts of positive selection-driven amino acid substitutions which we could not detect with certainty. This would be the case for any burst that occurred within an external edge of a tree, or within an internal edge that is not short enough, or even within a short internal edge as long as the burst itself involved only a small number of substitutions. Unfortunately, we cannot estimate the number of such real but not confidently detectable bursts.

Obviously, our ability to detect a burst depends on the length of the internal edge. Roughly speaking, all bursts that involve at least ~8 amino acid substitutions within a protein of <300 amino acids can be detected within internal edges of length below 0.005 dS. Because we investigated 3411 proteins and the total length of all such edges in the gammarid tree was 0.15 dS, 5 bursts that we found in them imply that during an interval of time required for 1 synonymous substitution to occur per site, a protein undergoes a strong burst of adaptive evolution with probability ~0.01 (the estimate derived from primates is similar). If so, such bursts are not uncommon.

What can we say about the genes that underwent bursts? Not much: they evolve faster than an average gene, but only marginally so. Unexpectedly, 3 out of 5 genes encode mitochondrial proteins: a mitochondrial chaperone and two mitochondrial ribosome proteins. This observation is hard to explain. Mitochondrial genomes of Baikal Lake gammarids have been shown to undergo intensive rearrangement, which in combination with mito-nuclear discordance and epistatic interactions between mitochondrial proteins coded in nuclear and mitochondrial genomes might lead to this phenomenon (Romanova *et al.*, 2016).

Correlated positive selection at mutiple sites that leads to a burst may emerge due to a variety of mechanisms. One possibility, of course, is a sudden, drastic change of the adaptive landscape of a protein. However, a burst can also occur as a result of only a small change of the landscape, if it caused a fold bifurcation which eliminated a fitness peak that was occupied by a protein (Dodson and Hallam, 1977, Fig. 4). This mechanism is compatible with the fact that all genes with bursts in gammarids show a low overall dN/dS ratio on the entire phylogenetic tree (<0.57), implying that these bursts of evolution affected genes that usually evolve slowly. By contrast, the PKR gene of primates possessed a high dN/dS ratio (>1), implying that the burst in this gene involved an episode of additional acceleration of evolution which was generally fast.

## Acknowledgements

This study was supported by Russian Science Foundation (grant 16-14-10173) to ASK.

## Author contributions

AVS carried out the data analyses, participated in the design of the study and drafted the manuscript; GAB participated in the design of the study and helped draft the manuscript; TVN provided experimental validation of the results; ASK conceived of the study and helped draft the manuscript. All authors gave final approval for publication.

